# Social play is critical for the development of prefrontal inhibitory synapses and cognitive flexibility in rats

**DOI:** 10.1101/2020.05.01.070540

**Authors:** Ate Bijlsma, Azar Omrani, Marcia Spoelder, Jeroen P.H. Verharen, Lisa Bauer, Cosette Cornelis, Beleke de Zwart, René van Dorland, Louk J.M.J. Vanderschuren, Corette J. Wierenga

## Abstract

Sensory driven activity during early life is critical for setting up the proper connectivity of the sensory cortices. Here we ask if social play behavior, a particular form of social interaction that is highly abundant during post-weaning development, is equally important for setting up connections in the developing prefrontal cortex (PFC). Young rats were deprived from social play with peers for 3 weeks during the period in life when social play behavior normally peaks (P21-42; SPD rats), followed by resocialization until adulthood. We recorded synaptic currents in L5 cells in slices from medial PFC of adult SPD and control rats and observed that inhibitory synaptic currents were reduced in SPD slices, while excitatory synaptic currents were unaffected. This was associated with a decrease in perisomatic inhibitory synapses from parvalbumin-positive GABAergic cells. In parallel experiments, adult SPD rats achieved more reversals in a probabilistic reversal learning task (PRL), which depends on the integrity of the PFC. They appeared to use a different cognitive strategy than controls. One hour of intense play during SPD did not prevent the decrease in inhibitory synaptic inputs and had only a limited effect on behavioral outcomes in the PRL. Our data demonstrate the importance of unrestricted social play for the development of inhibitory synapses in the PFC and cognitive skills in adulthood.

## Introduction

The developing brain requires proper external input to fine-tune activity and connectivity in neural circuits to ensure optimal functionality throughout life. This process has been extensively studied in the sensory cortices, and it is long known that sensory deprivation during development causes long-lasting deficits in sensory processing resulting from improper synaptic wiring (Hensch, 2005; Gainey and Feldman, 2017). However, how experience-dependent plasticity contributes to the development of other brain structures, such as the prefrontal cortex (PFC), remains largely unknown (Kolb et al., 2012; Larsen and Luna, 2018; Reh et al., 2020).

The PFC is important for higher cognitive, so-called executive functions (Miller and Cohen, 2001; Floresco et al., 2008), as well as the neural operations required during social interactions (Frith and Frith, 2012; Rilling and Sanfey, 2012). By analogy of sensory cortex development, proper PFC development may require complex cognitive and social stimuli. Importantly, during the period when the PFC matures, i.e. in between weaning and early adulthood (Kolb et al., 2012), young animals display an abundance of an energetic form of social behavior known as social play behavior (Panksepp et al., 1984; Vanderschuren et al., 1997; Pellis and Pellis, 2009). Social play behavior involves PFC activity (van Kerkhof et al., 2013b),and lesions or inactivation of the PFC have been found to impair social play (Bell et al., 2009; van Kerkhof et al., 2013a). It is widely held that exploration and experimentation during social play facilitates the development of a rich behavioral repertoire, that allows an individual to quickly adapt in a changeable world. In this way, social play subserves the development of PFC-dependent skills such as flexibility, creativity, and decision-making (Spinka et al., 2001; Pellis and Pellis, 2009; Vanderschuren and Trezza, 2014).

Lack of social play experience during post-weaning development may cause long-lasting changes in PFC circuitry and function (Leussis et al., 2008; Bell et al., 2010; Baarendse et al., 2013; Vanderschuren and Trezza, 2014). However, the cellular mechanisms by which social play facilitates PFC development remain elusive. It is well described that sensory deprivation induces specific alterations in inhibitory neurotransmission that affect adult sensory processing (Turrigiano and Nelson, 2004; Hensch, 2005). We therefore hypothesized that early life social experiences specifically shape PFC inhibition. Here we investigated how cognitive flexibility and inhibitory signaling are affected in the adult PFC when rats are deprived from social play during development.

## Results

Rats were deprived of social play for a period of 3 weeks post-weaning (postnatal days 21-42), which is the period in life when social play is most abundant (Baenninger, 1967; Meaney and Stewart, 1981; Panksepp, 1981) . Rats in the social play deprivation (SPD) group were separated from their cage mate by a plexiglass wall, which allowed smelling, hearing, seeing and communicating, but not physical interaction and playing. After the SPD period, the wall was removed and pair-wise social housing was maintained until adulthood when experiments were performed (postnatal week 8-10). Control (CTL) rats were housed in pairs during the entire period. We performed voltage-clamp recordings from layer 5 (L5) pyramidal cells of the medial PFC (mPFC) in slices prepared from adult SPD and CTL rats to assess the impact of SPD on PFC circuitry development (Fig. 1A-C). We found that the frequency and amplitude of spontaneous inhibitory postsynaptic currents (sIPSCs) was reduced in SPD rats (Fig 1D, E). By contrast, spontaneous excitatory postsynaptic currents (sEPSCs) were unaffected (Fig. 1F, G). The frequency of miniature inhibitory currents (mIPSCs) was also reduced in SPD slices (Fig. 1H), while mIPSC amplitudes (Fig. 1I) were not affected. The reduction in mIPSC frequency in SPD slices was accompanied by an increase in the average rise time (Fig. 1J), suggesting that particularly mIPSCs with fast kinetics were lost. Decay kinetics (Fig. 1K) and intrinsic excitability (Fig. 2A-C) were unaffected. Together, these data indicate that SPD leads to selective reduction in GABAergic synaptic inputs onto L5 pyramidal cells in the adult mPFC.

**Figure 1.**
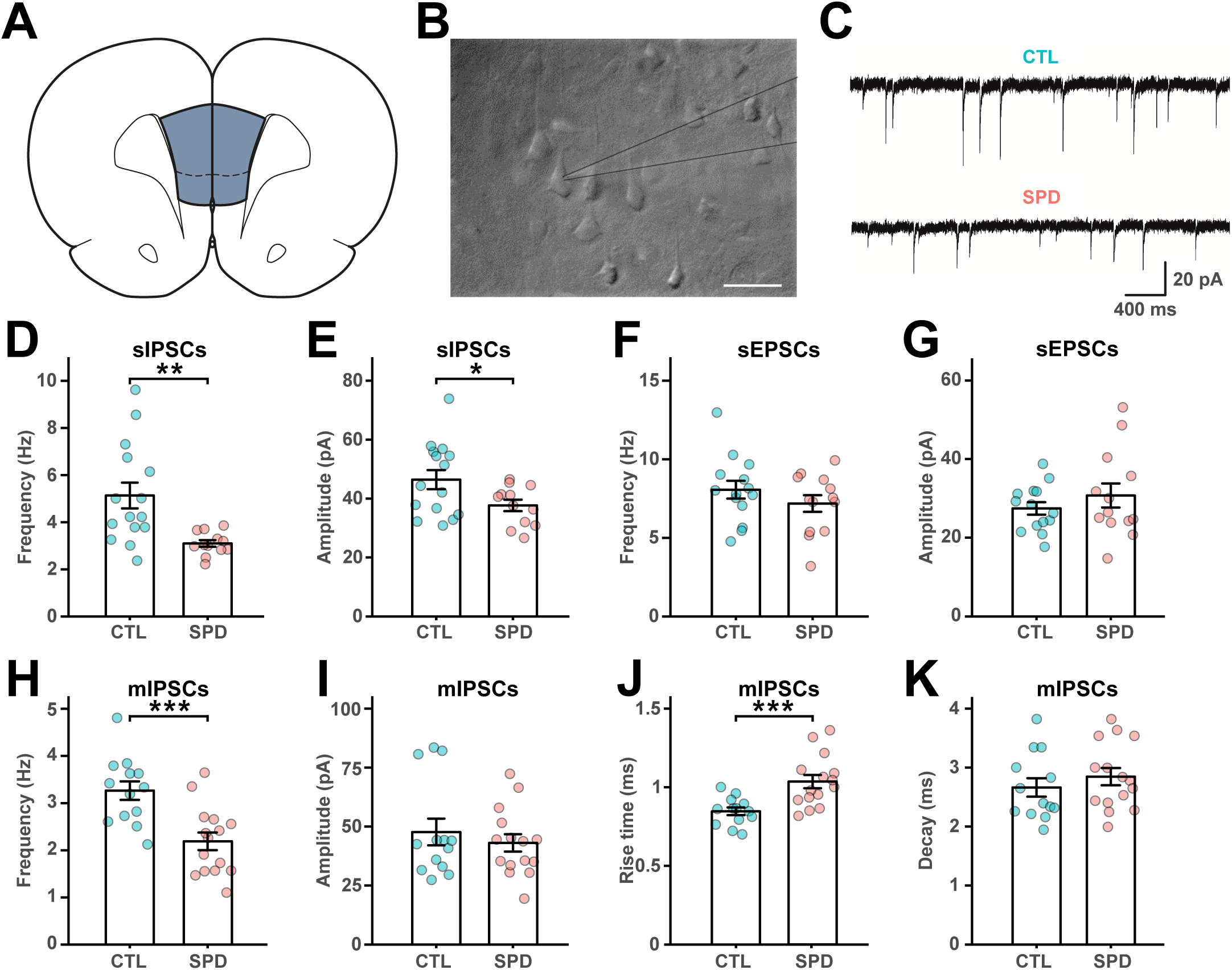
Reduced prefrontal inhibition in L5 pyramidal cells after social play deprivation. (A) Schematic diagram depicting the recording site in the mFPC. (B) DIC image of L5 cells in the mPFC with the recording electrode (grey lines). Scale bar is 20 μm. (C) Example traces of spontaneous IPSCs (sIPSCs) in L5 pyramidal cells in slices from control (CTL) and SPD rats. (D, E) Frequency (D) and amplitude (E) of sIPSCs in CTL and SPD slices (p = 0.002 and p = 0.03; T test). (F, G) Frequency (F) and amplitude (G) of spontaneous EPSCs (p = 0.27 and p = 0.35; T test). (H, I) Frequency (H) and amplitude (I) of miniature IPSCs (p < 0.0005 and p = 0.50; T test). (J) Rise time of mIPSCs (p = 0.0008; T test). (K) Decay time of mIPSCs (p = 0.40; T test). Data from 15 CTL and 13 SPD brain slices (6 rats per group). Statistical range: * p<0.05; ** p<0.01; *** p<0.001

**Figure 2.**
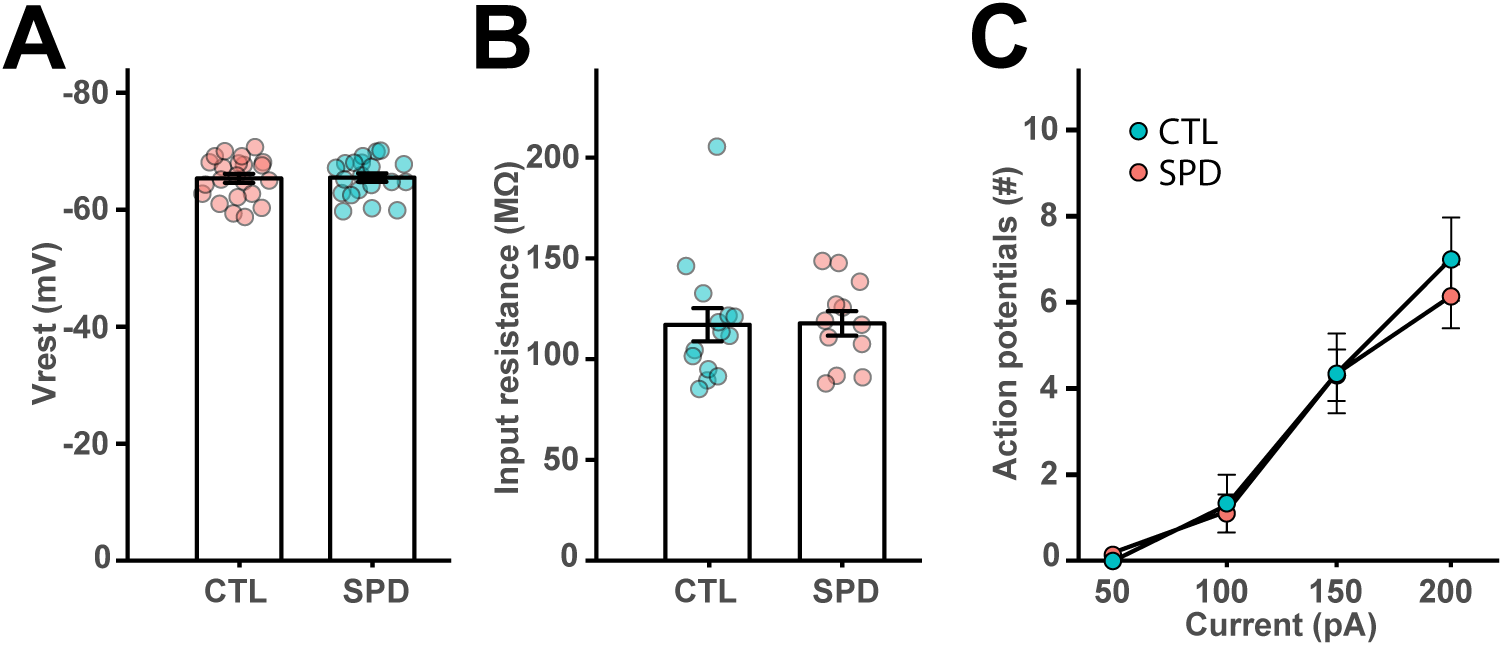
Passive membrane properties of L5 cells are similar in SPD and CTL slices. (A) Resting potential of L5 pyramidal neurons in CTL and SPD slices (p = 0.97; T test). (B) Input resistance (p = 0.61; MW test). (C) Number of action potentials after current injections in CTL and SPD neurons (p = 0.58; 2w ANOVA, condition). Data in A is from 20 CTL and 22 SPD cells; in B from 14 CTL and 12 SPD cells; in C from 19 CTL and 22 SPD cells.

A reduction in mIPSCs with fast rise times suggests that inhibitory synapses at perisomatic locations were affected. Perisomatic synapses are made by parvalbumin (PV) and cholecystokinin (CCK) basket cells (Whissell et al., 2015), of which only the latter express the cannabinoid receptor 1 (CB1-R) (Katona et al., 1999). We performed immunohistochemistry on the mPFC of adult SPD and CTL rats and quantified the number of GAD67- and PV-positive cell bodies (Fig. 3A). The density of GAD67-positive interneurons (Fig. 3B) and PV-positive cells (Fig. 3C) in the mPFC was not different between SPD and CTL rats. We then quantified inhibitory synaptic markers around the soma of L5 pyramidal neurons, using NeuN staining to draw a narrow band around the soma of individual pyramidal neurons and analyzing the synaptic PV and CB1 puncta within this band (Fig. 4A-B; see Methods for details). The density of VGAT puncta was not different in SPD and CTL tissue, but the density of PV synapses (colocalizing with VGAT) was significantly lower in SPD tissue compared to CTL tissue (Fig. 4C). The density of CB1-R synapses was not altered. In addition, synaptic PV and CB1-R puncta intensity was decreased in SPD tissue (Fig. 4D). This reduction was specific as VGAT puncta intensity was similar in SPD and CTL tissue (Fig. 4D). Puncta size was not much affected, although synaptic CB1-R puncta were slightly larger in SPD tissue (Fig. 4E). We verified that somata of L5 pyramidal cells had similar size in SPD and CTL tissue (Fig. 4F) and synaptic density was not correlated with cell size (Fig. 4G). Together, these data show that SPD results in a decrease in inhibitory synaptic input to L5 pyramidal cells in the adult PFC, with differential effects on perisomatic PV and CB1-R synapses.

**Figure 3.**
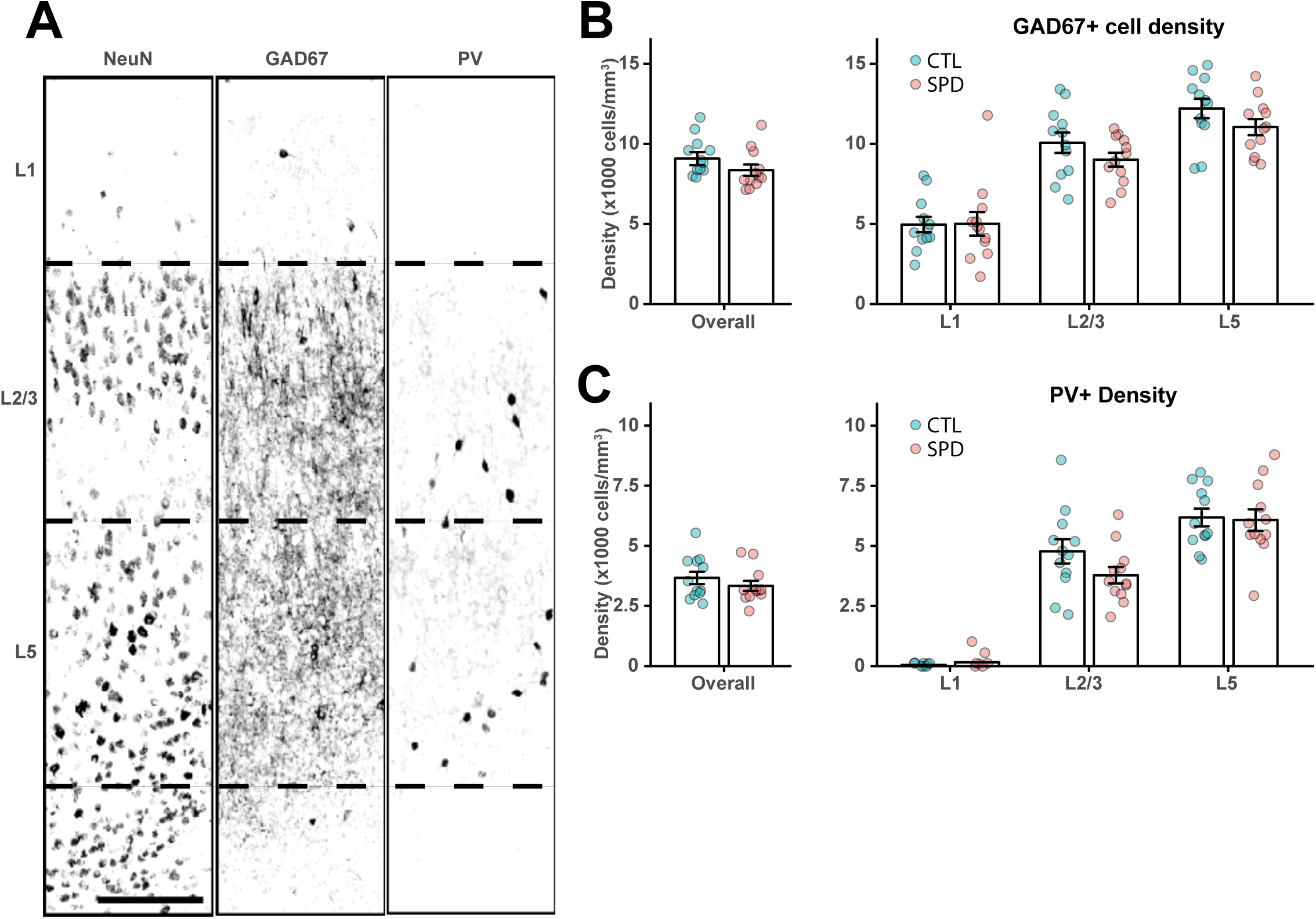
Interneuron density is similar in CTL and SPD tissue. (A) Representative confocal image of NeuN, GAD67 and PV positive neurons in prelimbic cortex layers. Borders between layers are denoted by the dashed lines. Scale bar is 10 μm. (B) Left: The average density of GAD67-positive cells in the mPFC over all layers (p = 0.19; T test); right: GAD67 cell density in Layer 1 (p = 0.95; T test), Layer 2/3 (p = 0.19; T test) and Layer 5 (p = 0.15; T test). (C) Left: The average density of PV-positive cells in the mPFC over all layers (p = 0.33; T test); right: PV cell density in Layer 1 (p = 0.90; MW test), Layer 2/3 (p = 0.12; T test) and Layer 5 (p = 0.85; T test). Data in B and C from 6 CTL and 6 SPD rats. For each rat two measurements (from both hemispheres) were included.

**Figure 4.**
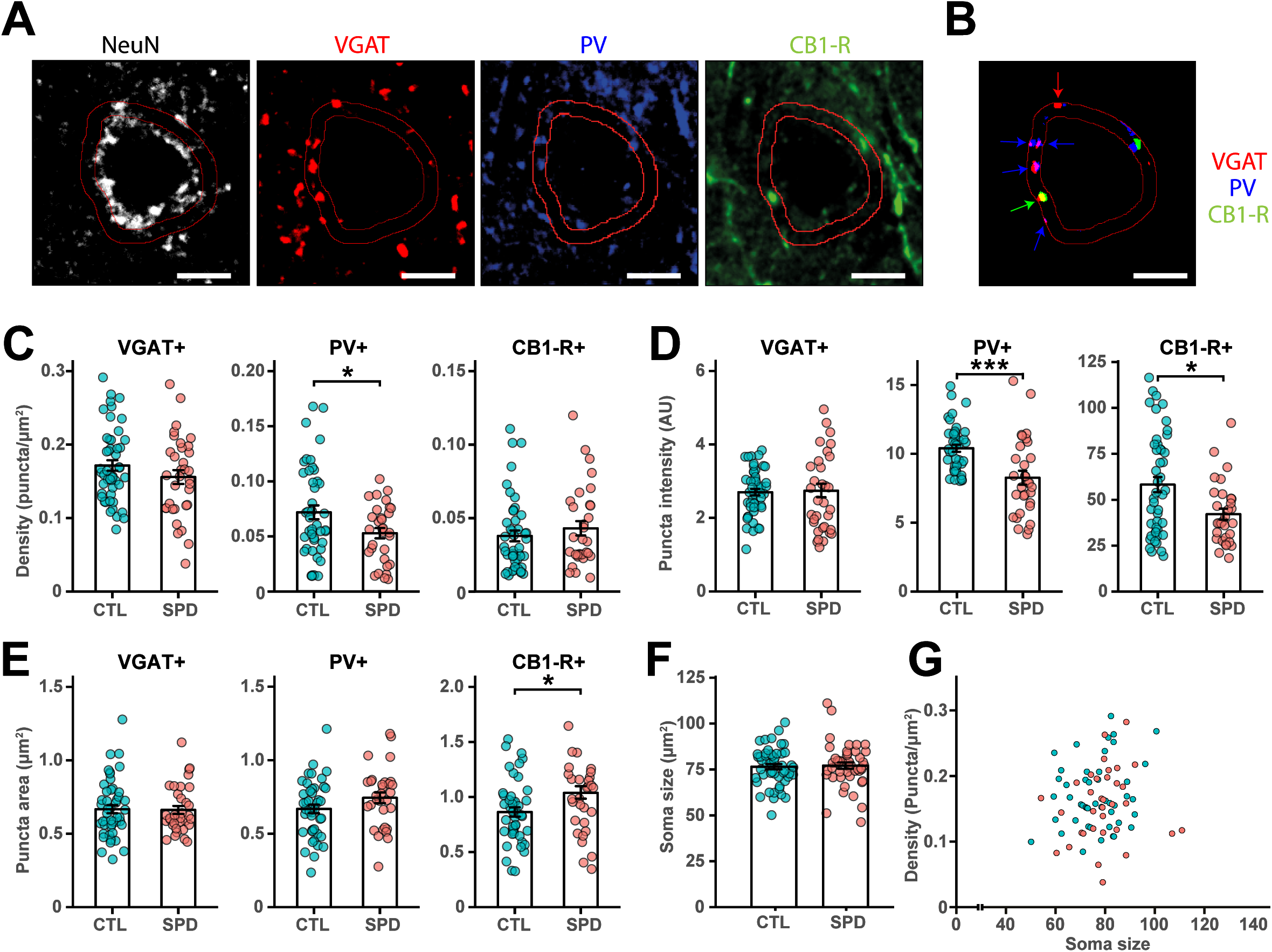
Reduction in perisomatic inhibitory synapses after SPD. (A, B) Representative confocal images for VGAT, PV, CB1-R and NeuN immunostaining. Scale bar is 1 μm. (C) A 1.5 μm band around the soma was drawn based on the NeuN staining. Individual puncta were selected after thresholding Only PV and CB1-R puncta that co-localized with VGAT were considered synaptic puncta. (D) Summary of the selected VGAT, PV and CB1-R puncta from C. (E) The density of synaptic VGAT, PV and CB1-R puncta (VGAT p = 0.19; PV p = 0.02; CB1-R p = 0.36) (F) The mean intensity of synaptic puncta (VGAT p = 0.51; PV p < 0.0005; CB1-R p = 0.02) (G) The mean area of synaptic puncta (VGAT p = 0.93; PV p = 0.12; CB1-R p = 0.05) (H) Soma size of L5 pyramidal cells (p = 0.20; T test). (I) Correlation between L5 soma size and VGAT synaptic puncta density. Data in E-I from 49 CTL cells and 34 SPD cells (6 rats per group, 2 hemispheres). Statistical range: * p<0.05; ** p<0.01; *** p<0.001.

In order to assess the impact of SPD on cognitive flexibility, a PFC-dependent probabilistic reversal learning task (PRL) was used (Fig. 5A, B) (Dalton et al., 2016; Verharen et al., 2020). In this task, responding on the ‘correct’ and ‘incorrect’ levers was rewarded on 80% and 20% of trials, respectively, and position of the ‘correct’ and ‘incorrect’ levers switched after 8 consecutive responses on the ‘correct’ lever. Rats in the SPD and CTL groups readily acquired the task and achieved a comparable performance level in terms of rewards obtained (Fig. 5C). Remarkably, SPD rats completed more reversals than CTL rats (Fig. 5D), indicating that cognitive performance in adult rats was altered after SPD.

**Figure 5.**
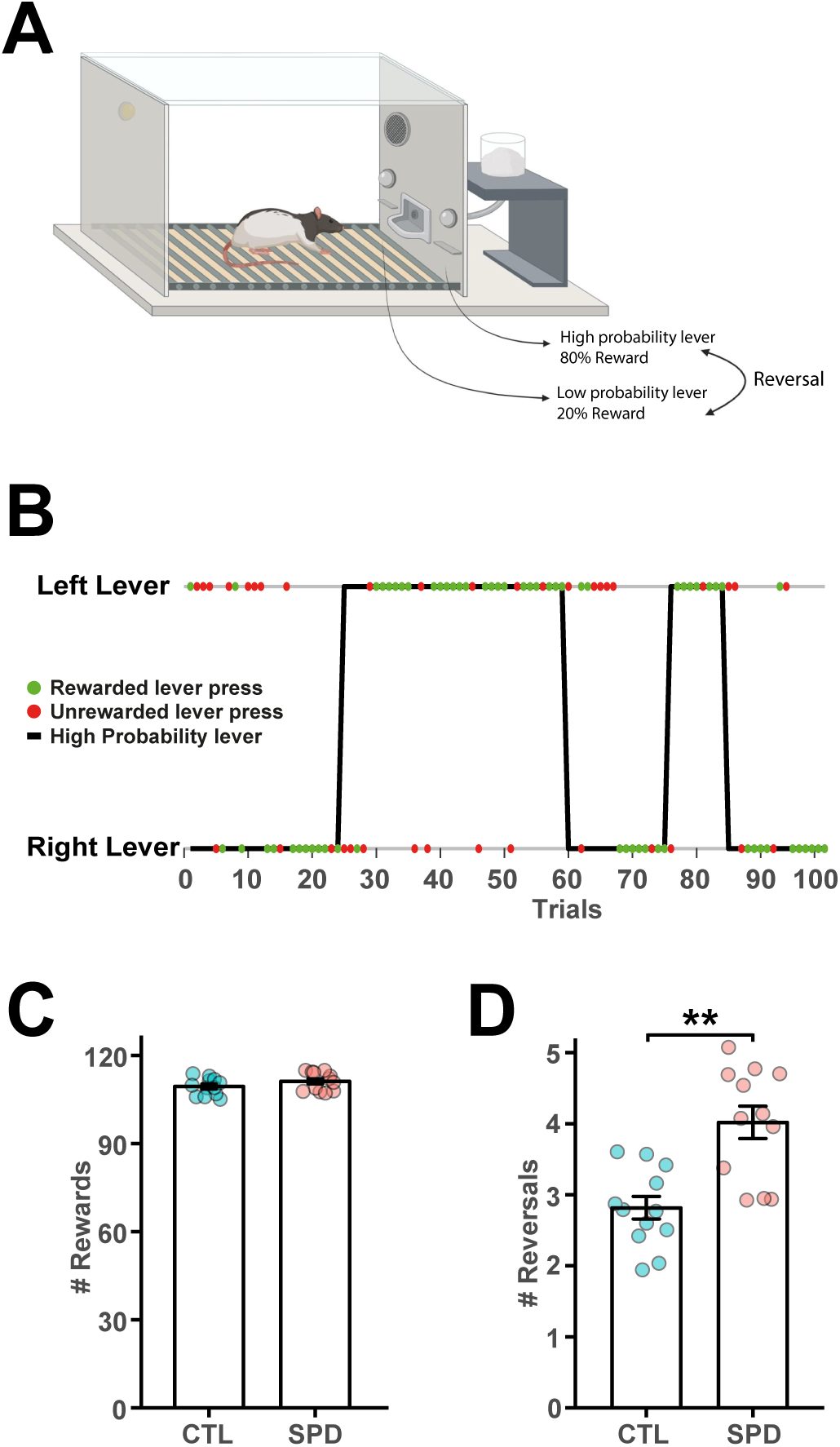
Altered PRL performance after SPD. (A) Probabilistic reversal learning task design. Reversals occur when the rat has pressed the high probability lever 8 consecutive times. (B) Representation of the first 100 lever presses of an example rat during one of the sessions. Green and red dots represent rewarded and unrewarded lever presses respectively. A reversal is indicated by the chance of the high probability lever (Trial 24, 59, 75 and 84). (C) Average number of rewards for CTL and SPD rats (p = 0.16; T test). (D) Average number of reversals (p = 0.001; T test). Data in C and D from 12 CTL and 12 SPD rats. Statistical range: ** p > 0.01.

Early behavioral studies have shown that behavioral performance after social isolation can be partially rescued by allowing the rats daily play for a short amount of time (Einon et al., 1978; Potegal and Einon, 1989). We therefore repeated the PRL experiments, but now added a second group of SPD rats which were allowed to play daily for 1 hour with their cage mate during the deprivation period (SPD1h). We quantified pinning and pouncing - the most characteristic social play behaviors in rats (Panksepp and Beatty, 1980; Vanderschuren et al., 1996) -, as well as social and non-social exploration during the play sessions of the SPD1h rats (Fig. 6A-D). The frequency of pins and pounces was much higher compared to socially housed rats of the same age, which typically show 1-2 pins per minute (Schneider et al., 2016; Stark and Pellis, 2021), and was comparable to rats in other studies that were isolated for 24 h, an isolation period that is known to induce maximal social play behavior (Niesink and Van Ree, 1989; Vanderschuren et al., 1995, 2008; Achterberg et al., 2016). Quantification of the social interactions showed that around 75 % of pins and pounces occurred in the first half hour of the session (Fig. 6E).

**Figure 6.**
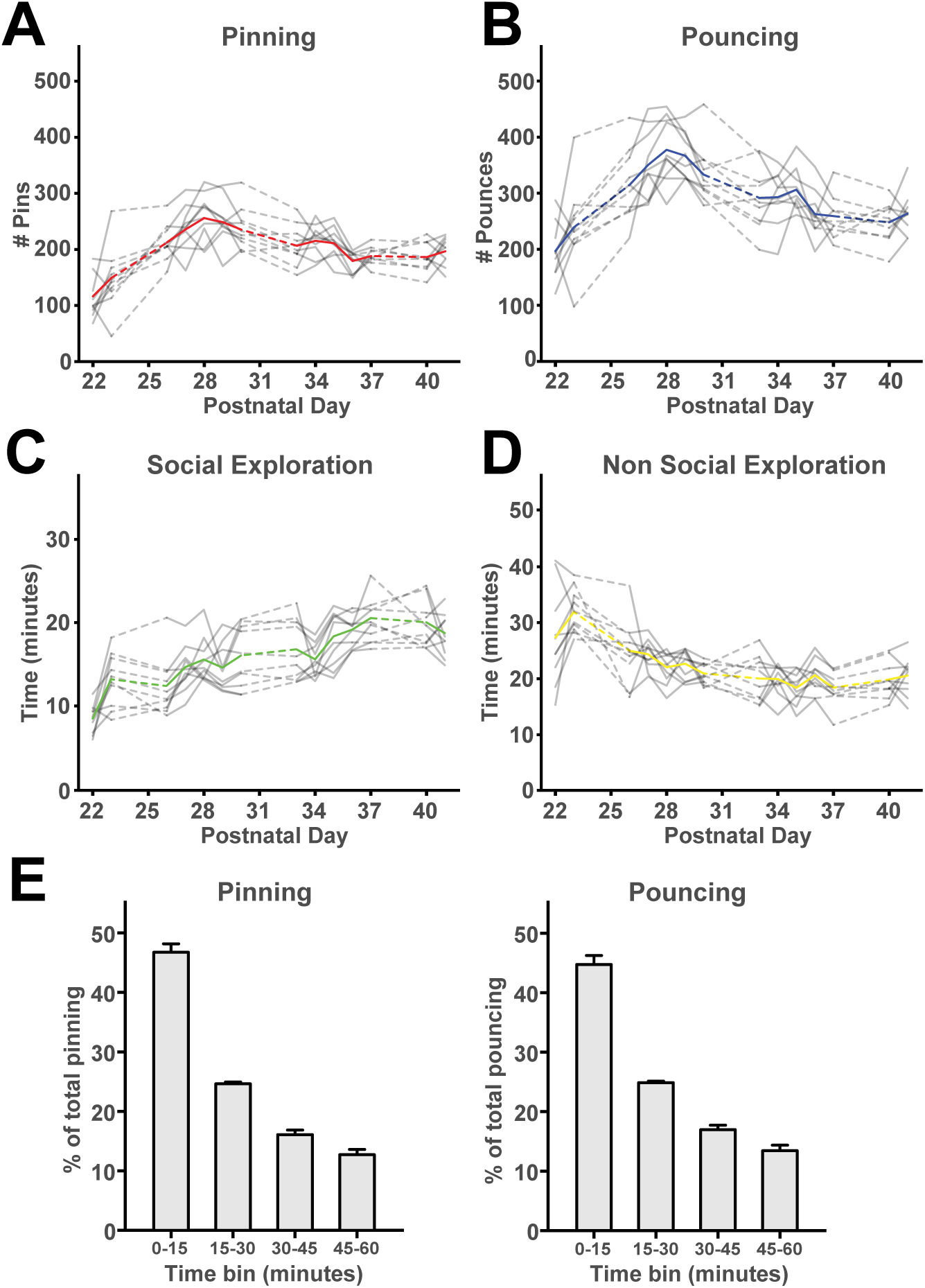
Social behavior during 1-hour play sessions. (A, B) The frequency of (A) pinning and (B) pouncing of the SPD1h rats during the 1-hour play sessions per day. (C, D) The time spent in (C) social and (D) non-social exploration. Each grey line represents a pair of rats with the colored line representing the groups’ average. (E, F) The behavioral readouts of social play are expressed as fraction of the total. For both (E) pinning and (F) pouncing, the amount was separated in bins of 15 mins (For statistics see statistical table 1). Data from 10 couples of SPD1h rats.

**Table 1.** Overview of computational models

| Model family | # | Model name | $n_f$ | Description |
| --- | --- | --- | --- | --- |
| Random choice | 1 | Random choice model | 0 | Animal choses randomly |
| Heuristics family | 2 | Win-Stay, Lose-Switch | 1 | Animal stays on the same lever after winning, moves away from the lever after a loss. |
|  | 3 | Win-Stay, Lose-Random | 1 | Animal stays on the same lever after winning, randomly picks a lever after a loss. |
|  | 4 | Random choice + stickiness | 2 | Animal chooses randomly but attributes value to the lastly chosen lever (Verharen et al., 2020). |
| Q learning family | 5 | Q learning, single learning rate for reward and punishment learning | 2 | Animal learns from previous decisions. Learning rate for positive and negative feedback are the same (Verharen et al., 2020). |
|  | 6 | Q learning, separate learning rates for reward and punishment learning | 3 | Animal learns from previous decisions. Learning rate for positive and negative feedback may differ (Verharen et al., 2020). |
|  | 7 | Model 6 + stickiness parameter | 4 | Animal learns from previous decisions. Learning rate for positive and negative feedback are separately calculated. Additionally, the animal attributes value to the lastly chosen lever (Verharen et al., 2020). |
|  | 8 | Rescorla-Wagner Pearce Hall | 4 | Animal learns from previous decisions. Learning rate for positive and negative feedback are the same. Additionally, the animal may learn better when task volatility is higher (e.g., after a reversal). |

Consistent with our first observations (Fig. 5), rats in all three groups equally learned to perform this task (Fig. 7A). They achieved a comparable performance level, with only small differences in the number of earned rewards (Fig. 7B). Again, SPD rats completed more reversals compared to the CTL rats, and this was observed consistently across all sessions (Fig. 7C). Reversal completion of the SPD1h group resembled the SPD group during the initial sessions of the task, but their performance in later sessions was more comparable to the control group (Fig. 7C). To understand these differences between the groups in more detail, we assessed win-stay and lose-shift behavior. We observed a specific increase in win-stay behavior in SPD rats, but not in the SPD1h group (Fig. 7D). When we assessed this behavior over the course of the sessions (Fig. 7E), we observed more win-stay choices in the SPD group compared to CTL rats consistently in all sessions. Intermediate behavior was observed in SPD1h rats, whereby their behavior resembled the SPD group in early sessions and the CTL group in later sessions. There was no difference between groups in their choices after non-rewarded trials (Fig. 7F, G).

**Figure 7.**
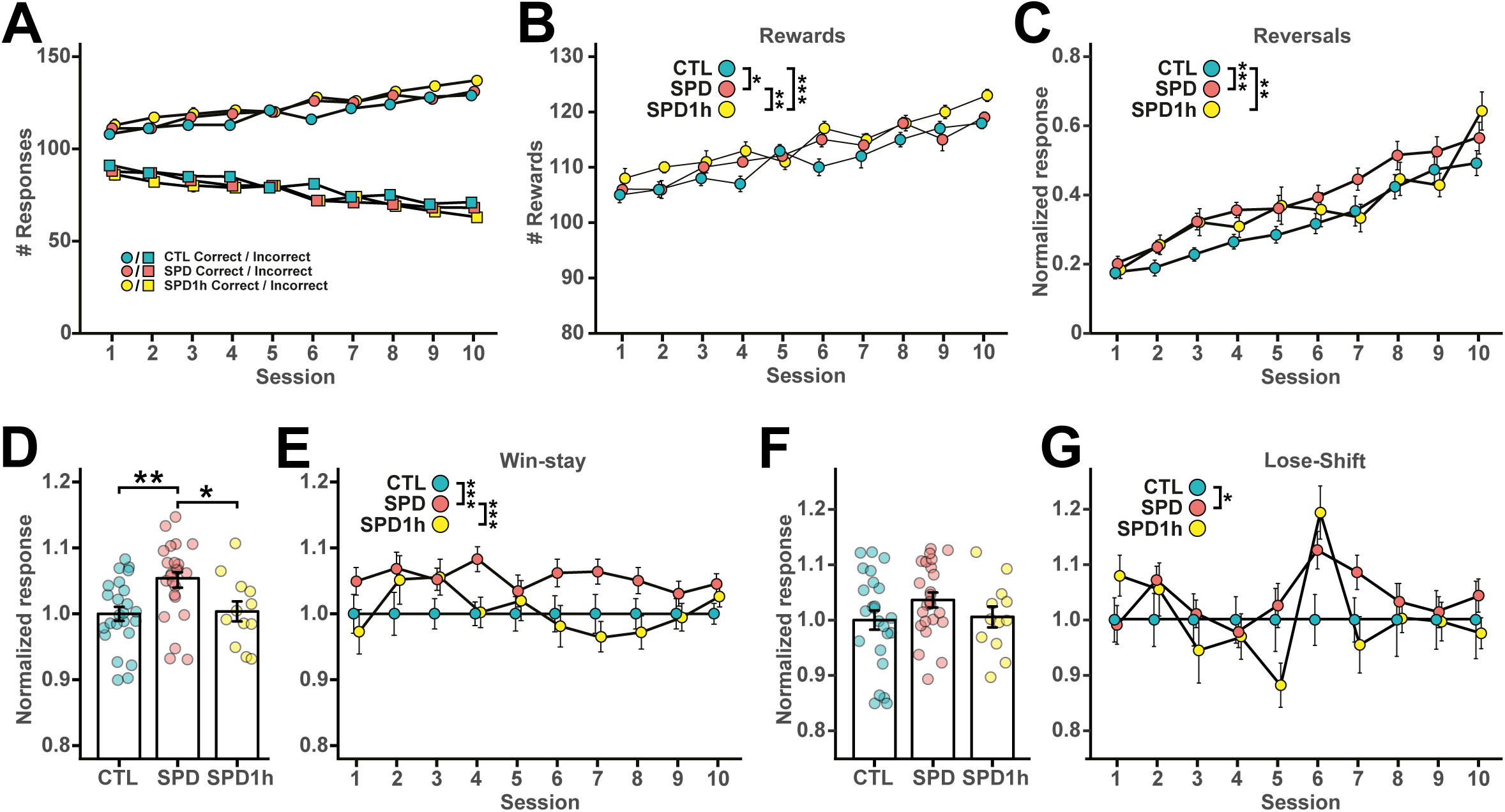
Behavioral analysis of altered PRL performance after SPD. (A) Number of correct and incorrect lever presses per session for CTL, SPD and SPD1h rats. (B) Number of sucrose rewards earned per session. (C) Normalized number of reversals per session. (D) Normalized average win-stay responses. (E) Normalized win-stay responses per session. (F) Normalized average lose-shift responses. (G) Normalized lose-shift responses per session. For statistics see statistical table 1. Data from 24 CTL, 24 SPD and 12 SPD1h rats (including CTL and SPD rats from fig. 5). Statistical range: * p<0.05; ** p<0.01; *** p<0.001

We next performed trial-by-trial analysis of the behavioral data (Verharen et al., 2018, 2020) to reveal possible alterations in the component processes subserving probabilistic reversal learning. We compared different computational models to describe the behavioral choices of the rats. The simplest random model assumes that animals always randomly choose a lever to press. A family of three heuristic models assumes simple practical strategies to complete the task (e.g. win-stay; lose-shift, etc). Finally, the four Q learning models integrate sensitivity to positive and negative feedback, and weigh exploration versus exploitation (see Methods for details). The behavior of CTL rats was best described by a Q-learning model in all sessions (Fig. 8A, B left panel), but the SPD and SPD1h rats shifted towards behavior congruent with a simpler heuristic strategy as reversal learning progressed (Fig. 8A, B middle and right panel), showing a tendency to remain at the previously chosen lever. Thus, SPD rats, with or without daily playtimes, switched from a learning-based strategy to a more perseverative strategy, whereas CTL rats continued learning throughout training. These data indicate that juvenile SPD alters PFC function and cognitive flexibility in adulthood. Furthermore, 1hr daily play during SPD only partially restores the cognitive performance in SPD rats.

**Figure 8.**
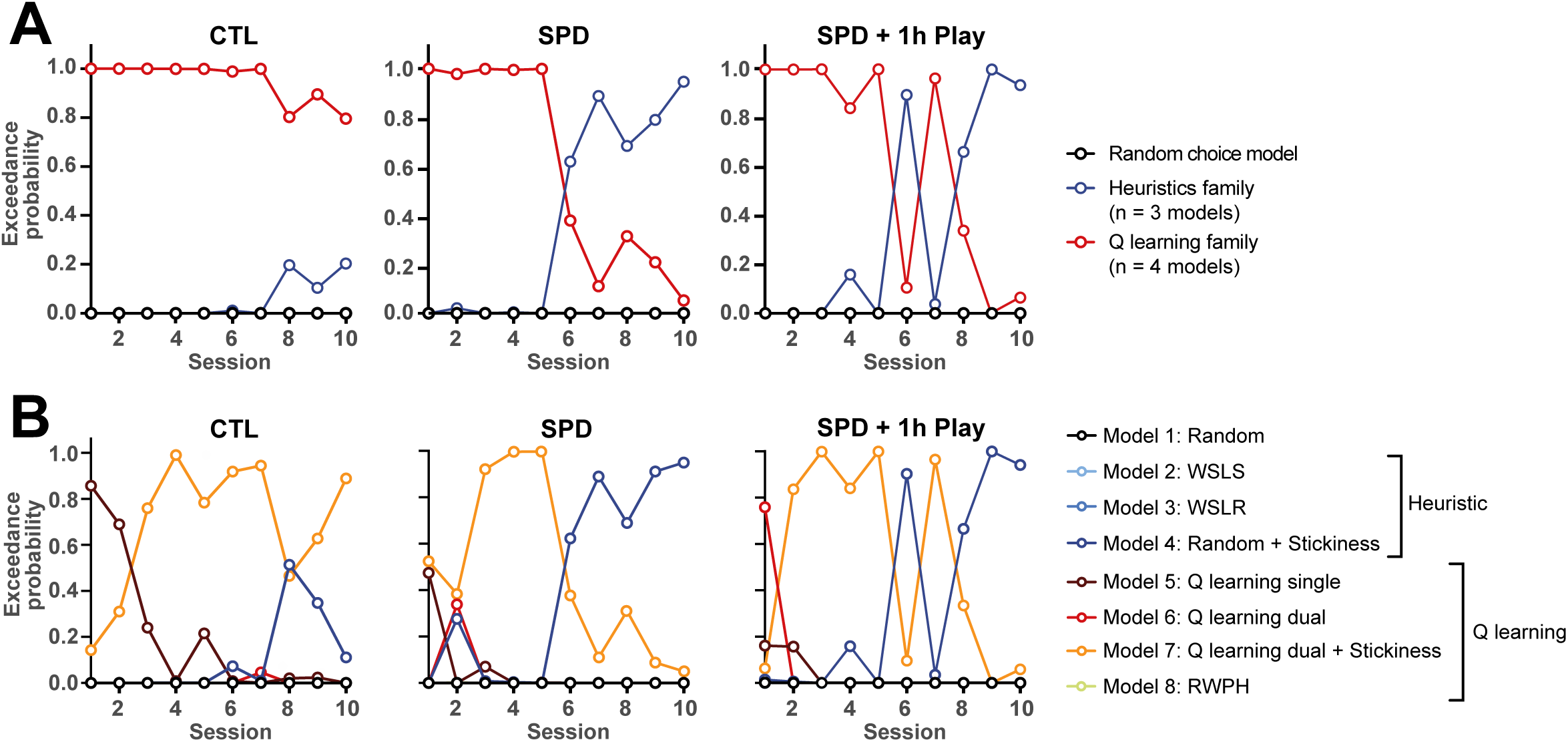
Trial-by-trial analysis of PRL performance. (A) Exceedance probability for different families of computational models (Random choice, Heuristics and Q-learning families) based on Bayesian model selection for the three groups (CTL, SPD and SPD1h rats). (B) Exceedance probability for random and specific heuristic (Win-stay/Lose-shift (WSLS), Win-stay/Lose-random (WSLR), Random + stickiness) and Q-learning family models (Q-Learning single, dual, dual + stickiness, Rescorla-Wagner-Pearce-Hall (RWPH)).

To check if the partial rescue in behavior of the SPD1h rats was due to a rescue of the inhibitory microcircuitry in the mPFC, we recorded miniature inhibitory currents (mIPSCs) in prefrontal slices from all 3 groups. Consistent with our earlier results (Fig 1H), a clear reduction in mIPSC frequency was found in the SPD group compared to CTL rats (Fig. 9B). This reduction was similar in the SPD and SPD1h slices, indicating that daily playtimes during the play deprivation period did not affect the development of PFC inhibition (Fig. 9B). Amplitudes of the events were not different between groups (Fig. 9C).

**Figure 9.**
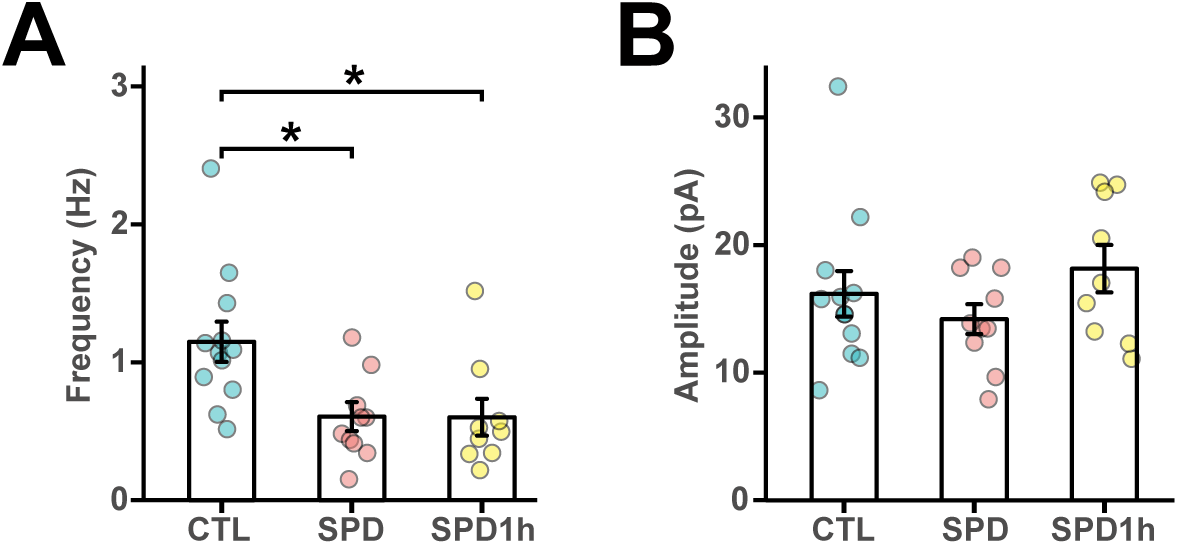
Reduced prefrontal inhibition in L5 pyramidal cells in SPD and SPD1h slices. (A) Frequency of mIPSCs in CTL, SPD and SPD1h slices (ANOVA p=0.0049, Tukey’s: CTL vs SPD p = 0.011; CTL vs SPD1h p = 0.017; SPD vs SPD1h p = 0.99). (B) Amplitude of mIPSCs in CTL, SPD and SPD1h slices (ANOVA p = 0.29). Data from 12 CTL, 10 SPD and 9 SPD1h cells (6 rats per group). Statistical range: * p < 0.05.

Our findings demonstrate that the organization of the adult GABAergic system in the mPFC of rats is robustly altered when rats are deprived of social play behavior during development, with important consequences for cognitive flexibility. The inclusion of daily play sessions during the SPD period failed to rescue the reduction in PFC GABAergic synapses and only partially restored the cognitive performance in a PFC-dependent PRL task.

## Discussion

When they are young, rats, like most other mammalian species, display an abundance of a particular, highly rewarding and energetic form of social behavior, termed social play behavior (Panksepp et al., 1984; Vanderschuren et al., 2016; Pellis and Pellis, 2017). Playing with peers it thought to allow young animals to experiment with their behavioral repertoire, and to provide practice scenarios to obtain the social, cognitive and emotional skills to become capable adults who can easily navigate a changeable world (Spinka et al., 2001; Pellis and Pellis, 2009; Vanderschuren and Trezza, 2014; Larsen and Luna, 2018). Social play enhances neuronal activity in a broad network of limbic and corticostriatal structures (Gordon et al., 2002, 2003; van Kerkhof et al., 2013b). This integrated neuronal activity is likely to induce synaptic plasticity in the PFC, analogous to the well-described experience-dependent maturation of cortical sensory circuits, which requires appropriate sensory activation during development (Hensch, 2005; Gainey and Feldman, 2017). Our study investigates at the synaptic level how social play experience in juvenile rats – roughly equivalent to childhood in humans - contributes to maturation of PFC circuitry. Our data show that the experience of unrestricted juvenile social play is crucial to instruct the development of specific inhibitory connections in the PFC and to shape adaptive cognitive strategies in the adult brain.

We found that perisomatic inhibition onto L5 cells was reduced in the adult mPFC after SPD. We observed a ∼30% reduction in mIPSC frequency, associated with a comparable reduction in perisomatic PV synapses. A preferential effect of SPD on perisomatic inhibition by PV-expressing cells parallels observations after sensory deprivation during development (Hensch et al., 1998; Jiao et al., 2006; Mowery et al., 2019; Reh et al., 2020). The reduced levels of PV and CB1 expression that we observed suggest reduced PV activity (Donato et al., 2013; Caballero et al., 2014) and altered endocannabinoid tone (Sciolino et al., 2010; Schneider et al., 2016) in the PFC after SPD. This may interfere with developmental plasticity of PFC circuitry (Caballero and Tseng, 2016) and affect cognitive capacities in adulthood (Donato et al., 2015). Importantly and resonating well with our present findings, a reduction in PFC inhibition has previously been linked to impaired cognitive flexibility (Gruber et al., 2010), and recently a direct link between PFC PV cell activity and social behavior was demonstrated (Bicks et al., 2020; Sun et al., 2020).

Our data reveal specific synaptic alterations in the prefrontal microcircuitry that may underlie altered cognitive strategies in animals deprived of juvenile social play. Importantly, after the temporary social isolation when they were young, the animals had ample opportunity for social interaction for several weeks before testing. However, even after weeks of unrestricted social interactions, pronounced changes in PFC function and cognition persisted. This emphasizes the importance of early post-weaning social play, consistent with earlier studies that identified this time window as a critical period for PFC maturation and behavioral development (Einon and Morgan, 1977; Hol et al., 1999; Lukkes et al., 2009; Kolb et al., 2012; Whitaker et al., 2013).

Adult rats which had experienced SPD displayed a different behavioral strategy in the probabilistic reversal learning task compared to CTL animals. CTL rats seemed to follow a sophisticated cognitive strategy which was well-described by a Q-learning model. Our findings indicate that SPD rats performed more reversals – but actually received the same number of rewards compared to CTL rats – by using a simplified win-stay strategy, in which they relied less on feedback-driven learning, and more on perseveration-based heuristics. Such heuristics are thought to be cognitively less demanding (Christie and Schrater, 2015), and may therefore be preferred under certain conditions, especially if this does not lead to a reduction in reward. A similar increase in reversals was also reported after prelimbic mPFC inactivation (Dalton et al., 2016), suggesting that the mPFC may be less involved in PRL performance in SPD rats compared to CTL rats.

We observed that one hour of daily playtime during the deprivation period could not rescue the reduction in prefrontal IPSCs, and only partially restored behavior. Play is not displayed continuously by young rats, but appears in peaks of relatively short duration across the day (Melotti et al., 2014; Lampe et al., 2019). We observed that SPD1h rats played very intensely during the first 15-30 minutes of each play session resulting in a total amount of play that makes up a substantial fraction of the daily play that socially housed CTL rats show at this age (Baenninger, 1967; Vanderschuren et al., 2008; Schneider et al., 2016; Pellis and Pellis, 2017). However, it was not enough to prevent the reduction in prefrontal inhibition. This suggests the necessity of unrestricted, voluntarily elicited or repeated play such that short daily periods of intensive play cannot rescue the lasting effect of SPD on prefrontal inhibition and function. However, 1 hour of daily play partially rescued PRL performance. PRL performance depends on complex interactions between several component processes, including the sensitivity to positive and negative feedback, response persistence and exploration versus exploitation, each of which requires functional activity in distinct PFC regions (Verharen et al., 2018, 2020). Our observation that the behavior of SPD1hr fell in between SPD and CTL rats, while their reduction in mPFC inhibition was similar to SPD rats, suggests that the partial rescue of behavior involves compensatory changes in other brain areas, but further studies will be needed to elucidate this.

Together, our results demonstrate a key role for unrestricted juvenile social play in the development of perisomatic inhibition in the PFC, and PFC-dependent cognitive flexibility.

## Acknowledgements

Supported by the Netherlands Organisation for Scientific Research (NWO) ALWOP.2015.105 (LJMJV, CJW) and UU strategic theme Dynamics of Youth. We thank Daniëlle Counotte for preliminary experiments and Ruth Damsteegt for practical assistance.

## Author Contribution

The study was conceived by CW and LV. Experiments were designed by AB, AO, MS, LV and CW. Experiments were performed by AB, AO, MS, LB, RD, CC and BdW. Analysis was performed by AB, AO, MS, JV, LB, BdW and CC. Manuscript was written by AB, LV and CW.

## Methods

### Animals and housing conditions

All experimental procedures were approved by the Animal Ethics Committee of Utrecht University and the Dutch Central Animal Testing Committee and were conducted in accordance with Dutch (Wet op de Dierproeven, 1996; Herziene Wet op de Dierproeven, 2014) and European legislation (Guideline 86/609/EEC; Directive 2010/63/EU). Male Lister Hooded rats were obtained from Charles River (Germany) on postnatal day (P) 14 in litters with nursing mothers. All rats were subject to a reversed 12:12h light-dark cycle with ad libitum access to water and food. Rats were weaned on P21 and were either subjected to one of the social play deprivation (SPD and SPD1h) groups or the control (CTL) group. Control (CTL) rats were housed in pairs during the entire period. SPD rats were pair-housed with a rat from a different mother. During P21 to P42 a transparent Plexiglas divider containing small holes was placed in the middle of their home cage creating two separate but identical compartments. SPD rats were therefore able to see, smell and hear one another but they were unable to physically socially engage. SPD1h animals were housed similarly to the SPD group but were allowed to socially interact with their cage mate for 1 hour per day. Social interaction sessions took place in a Plexiglas arena of 40 x 40 x 60 cm (l × w × h) with approximately 2 cm of wood shavings. The social interaction sessions were recorded and the behavior was manually scored per pair using the Observer XT 15 software (Noldus Information Technology BV, Wageningen, The Netherlands). Four behaviors were scored:

- Frequency of pinning: one animal lying with its dorsal surface on the floor with the other animal standing over it.
- Frequency of pouncing: one animal attempts to nose/rub the nape of the neck of the partner.
- Time spent in social exploration: one animal sniffing or grooming any part of the partner’s body.
- Time spent in non-social exploration: the animals exploring the cage or walk around.

A total of 10 pairs were recorded during their social interaction sessions. Four of these were eventually used for the electrophysiology experiments while the other six couples were used for behavioral testing. On P42, the Plexiglas divider was removed and SPD and SPD1h rats were housed in pairs for the remainder of the experiment. All rats were housed in pairs for at least 4 weeks until early adulthood (10 weeks of age) after which experimentation began. All experiments were conducted during the active phase of the animals (10:00 - 17:00). One week before the start of behavioral testing, the rats were subjected to food-restriction and were maintained at 85% of their free-feeding weight for the duration of the behavioral experiment. Rats were provided with ∼ 20 sucrose pellets (45mg, BioServ) in their home cage for two subsequent days before their first exposure to the operant conditioning chamber to reduce potential food neophobia. Rats were weighed and handled at least once a week throughout the course of the experiment.

### Probabilistic reversal learning

#### Apparatus

Behavioral testing was conducted in operant conditioning chambers (Med Associates, USA) enclosed in sound-attenuating cubicles equipped with a ventilation fan. Two retractable levers were located on either side of a central food magazine into which sugar pellets could be delivered via a dispenser. A LED cue light was located above each retractable lever. A white house light was mounted in the top-center of the wall opposite the levers. Online control of the apparatus and data collection was performed using MED-PC (Med Associates) software.

#### Pre-training

Rats were first habituated once to the operant chamber for 30 min in which the house light was illuminated and 50 sucrose rewards were randomly delivered into the magazine with an average inter-trial interval of 15 s between reward deliveries. On the subsequent days, the rats were trained for 30 min under a Fixed-Ratio 1 (FR1) schedule of reinforcement for a minimum of three consecutive sessions. The session started with the illumination of the house light and the insertion of both levers, which remained inserted for the remainder of the session. One of the two levers was the ‘correct’ lever rendering a reward when pressed, whereas pressing the other lever had no consequences. There was no limit other than time on the amount of times a rat could press the ‘correct’ lever. If the rat obtained 50 or more rewards in a session it was required to press the other lever the following day. If it obtained less than 50 rewards the rat was tested on the same schedule the next day. A trial started with an inter-trial-interval (ITI) of 5 s with the chamber in darkness, followed by the illumination of the house-light and the insertion of one of the two levers into the chamber. A response within 30 s on the inserted lever resulted in the delivery of a reward. If the rat failed to respond on the lever within 30 s, the lever retracted and the trial was scored as an omission. Rats were trained for ∼ 3-4 days to a criterion of at least 50 rewards and had to perform a lever press in more than 80% of the trials before progressing to the probabilistic reversal learning.

#### Probabilistic reversal learning

The protocol used for this task was modified from those of previous studies (Bari et al., 2010; Dalton et al., 2016; Verharen et al., 2020). At the start of each session one of the two levers was randomly selected to be ‘correct’ and the other ‘incorrect’. A response on the ‘correct’ lever resulted in the delivery of a reward on 80% of the trials, whereas a response on the ‘incorrect’ lever was reinforced on 20% of trials. Each trial started with a 5 s ITI, followed by the illumination of the house-light and the insertion of both levers into the chamber. After a ‘correct’ response, both levers retracted but the house light remained illuminated. In case the rat was rewarded, the house light remained illuminated, whereas the house light extinguished in case the rat was not rewarded on the ‘correct’ lever. An ‘incorrect’ response or a failure to respond within 30 s after lever insertion (i.e. omission) lead to the retraction of both levers, extinction of the house light so that the chamber returned to its ITI state. When the rat made a string of 8 consecutive trials on the ‘correct’ lever (regardless of whether they were rewarded or not), contingencies were reversed, meaning the ‘correct’ lever became the ‘incorrect’ lever and the previously ‘incorrect’ lever became the ‘correct’ lever. This pattern repeated over the course of a daily session. Daily sessions were completed upon performing 200 trials or after 60 minutes have passed, whichever occurred first.

#### Trial-by-trial analysis

This analysis was performed to assess the shifts in choice behavior between subsequent trials, in order to investigate the sensitivity to positive and negative feedback. Depending on whether the rat received a reward or not, it can press the same lever on the subsequent trial or shift towards the other lever, resulting in 4 different choices (i.e. win-stay, win-shift, lose-stay, lose-shift) for both the ‘correct’ and ‘incorrect’ lever. We calculated these choices (win-stay vs win-shift and lose-stay vs lose-shift) as percentages per session.

#### Normalization

The PRL task was performed twice with two different batches of animals. The first batch consisted of 12 CTL and 12 SPD animals. The second batch consisted of three groups of 12 rats (CTL, SPD and SPD1h). For the number of reversals we used the following normalization using the minimum and maximum values (*groupmin*, *groupmax*) per group:

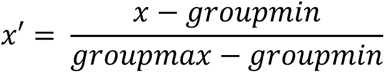

Where x is the original induvial value and x’ is the normalized value. To compare the win-stay and lose-shift choices per session, we normalized the SPD and SPD1h data to the average of their respective control group.

### Computational Modelling

Eight different computational models were fit to the trial-by-trial responses to assess differences in task strategy between the three groups of animals. Best-fit model parameters were estimated using maximum likelihood estimation, using Matlab (version 2018b; The MathWorks Inc.) function ‘fmincon’ (Verharen et al., 2018). These maximum likelihood estimates were corrected for model complexity (i.e., the number of free parameters (nf)) by calculating the Akaike information criterion (AIC) for each session:

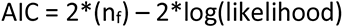

In which a lower AIC indicates more evidence in favor of the model. These log-model evidence estimates were subsequently used to perform Bayesian model selection (Rigoux et al., 2014) using the Matlab package SPM12 (The Wellcome Centre for Human Neuroimaging), taking into account the family to which each model belonged(Penny et al., 2010). This yielded the protected exceedance probability for the 8 individual models and for each family of models (random choice, heuristic and Q learning family), indicating the probability that each of the (family of) models was most prevalent among the group of rats.

Table 1 contains an overview of the eight computational models. The random choice model is the null model, which assumes that animals choose randomly (i.e., p = 0.5 for each choice, so that the log likelihood is given by [number of trials]*log(0.5)). The second family of models contained strategies based on ‘heuristics’; simple strategies to complete the task. The third family contained Q learning models, consisting of four derivatives of the Rescorla-Wagner model (Rescorla and Wagner, 1972).

### Electrophysiological analysis

The electrophysiology experiments were performed twice with two batches of different animals. The first batch consisted of 12 CTL and 12 SPD animals. The second batch consisted of three groups of 12 (CTL, SPD and SPD1h). *Slice preparation*: Rats (12-15 weeks of age) were anesthetized by intraperitoneal injection of sodium pentobarbital (batch 1) or induction with isoflurane (batch 2) and then transcardially perfused with ice-cold modified artificial cerebrospinal fluid (ACSF) containing (in mM): 92 N-methyl-D-glutamine (NMDG), 2.5 KCl, 1.25 NaH_2_PO_4_, 30 NaHCO_3_, 20 HEPES, 25 glucose, 2 thiourea, 5 Na-ascorbate, 3 Na-pyruvate, 0.5 CaCl_2_.4H_2_O, and 10 MgSO_4_.7H_2_O, bubbled with 95% O_2_ and 5% CO_2_ (pH 7.3–7.4). For batch 2 NMDG was replaced by choline chloride and thiourea was left out. Coronal slices of the medial PFC (300 µm) were prepared using a vibratome (Leica VT1200S, Leica Microsystems) in ice-cold modified ACSF. Slices were initially incubated in the carbogenated modified ACSF for 5-10 min at 34 °C and then transferred into a holding chamber containing standard ACSF containing (in mM): 126 NaCl, 3 KCl, 2 MgSO_4_, 2 CaCl_2_, 10 glucose, 1.25 NaH_2_PO_4_ and 26 NaHCO_3_ bubbled with 95% O_2_ and 5% CO_2_ (pH 7.3) at room temperature for at least 30 minutes (2 MgSO_4_ was replaced by 1.3 MgCl2 in batch 2). They were subsequently transferred to the recording chamber, perfused with standard ACSF that is continuously bubbled with 95% O_2_ and 5% CO_2_ at 28–32° C.

#### Whole-cell recordings and analysis

Whole-cell patch-clamp recordings were performed from layer 5 pyramidal neurons in the medial PFC. These neurons were visualized with an Olympus BX61W1 microscope using infrared video microscopy and differential interference contrast (DIC) optics. Patch electrodes were pulled from borosilicate glass capillaries and had a resistance of 3-6 MΩ when filled with intracellular solutions. Excitatory postsynaptic currents (EPSCs) were recorded in the presence of bicuculline (10 µM) and with internal solution containing (in mM): 140 K-gluconate, 1 KCl, 10 HEPES, 0.5 EGTA, 4 MgATP, 0.4 Na2GTP, 4 Na2phosphocreatine (pH 7.3 with KOH). Inhibitory postsynaptic currents (IPSCs) were recorded in the presence of 6-cyano-7-nitroquinoxaline-2,3-dione (CNQX) (10 µM in batch 1; 20 μM in batch 2) and D,L-2-amino-5-phosphopentanoic acid (D,L-AP5) (20 µM in batch 1; 50 μM in batch 2), with internal solution containing (in mM): 125 CsCl, 2 MgCl2, 5 NaCl, 10 HEPES, 0.2 EGTA, 4 MgATP, 0.4 Na2GTP (pH 7.3 with CsOH; batch 1) or 70 K-gluconate, 70 KCl, 10 HEPES, 0.5 EGTA, 4 MgATP, 0.4 Na2GTP, 4 Na2phosphocreatine (pH 7.3 with KOH; batch 2). Action-potential independent miniature IPSCs (mIPSCs) were recorded under the same conditions, but in the presence of 1 μM tetrodotoxin (TTX; Sigma) to block sodium channels. The membrane potential was held at -70 mV for voltage-clamp experiments. Signals were amplified, filtered at 3 kHz and digitized at 10 kHz using an EPC-10 patch-clamp amplifier with PatchMaster v2x73 software (batch 1) or MultiClamp 700B amplifier (Molecular Devices) with pClamp 10 software (batch 2). Series resistance was constantly monitored, and the cells were rejected from analysis if the resistance changed by >20%. No series resistance compensation was used. Resting membrane potential was measured in bridge mode (I=0) immediately after obtaining whole-cell access. The basic electrophysiological properties of the cells were determined from the voltage responses to a series of hyperpolarizing and depolarizing square current pulses. Input resistance was determined by the slope of the linear regression line through the voltage-current curve.

Passive and active membrane properties were analyzed with Clampfit 10 (Axon Instrument) or Matlab (Mathworks). Miniature and spontaneous synaptic currents (IPSCs and EPSCs) data were analyzed with Mini Analysis (Synaptosoft Inc., Decatur, GA). All events were detected with a criterion of a threshold >3× root-mean-square (RMS) of baseline noise. The detected currents were manually inspected to exclude false events.

### Immunohistochemistry

#### Tissue preparation

Rats were anesthetized with Nembutal (i.p. 240mg/kg) and transcardially perfused with 0.1 M phosphatebuffered saline (PBS, pH 7.3-7.4) followed by 4% paraformaldehyde in 0.01 M PBS. The brains were removed from the skull and post-fixed overnight in the same paraformaldehyde solution at 4◦C and subsequently cryoprotected in 30% sucrose for three days at 4◦C. Thereafter, the brains were rapidly frozen in aluminum foil on dry ice and stored at −80◦C until further use. Brain sections (20 μm thick) from the PFC between Bregma levels of 4.2 - 2.2mm were made with a Cryostat Leica CM 3050 S. Sections were stored at -80°C until immunohistochemistry was performed. Brain slices were thawed and let dry for 1h at room temperature (RT). Slices were washed in PBS three times for 15 min (3x15min) at RT. Sections were cooked in sodium citric acid buffer (SCAB, 10mM sodium citric acid in demi water, pH6) for 10 min at 97°C in a temperature controlled microwave, cooled for 30 min at 4°C and washed again (3x15min in PBS). Slices were blocked with 400 µl of blocking buffer (10% normal goat-serum, 0,2% triton-X 100 in PBS) for 2h in a wet chamber at RT. Slices were incubated overnight at 4°C in the wet chamber with 250 µl of primary antibodies in blocking buffer. Sections were washed (3x15min in PBS), followed by incubation with the secondary antibodies in blocking buffer for 2h at RT in a wet chamber. After another wash step (3x15min in PBS), slides were mounted and stored at 4°C until image acquisition. Primary and secondary antibodies are listed in Table 2 (for cell density analysis) and Table 3 (for synaptic puncta analysis).

**Table 2A.** Primary antibodies for interneuron analysis

| Host | Epitope | Concentration | Company | Order number |
| --- | --- | --- | --- | --- |
| Rat | Ctip2 | 1:1000 | Abcam | ab18465 |
| Rabbit | PV | 1:1000 | Life technologies | PA1933 |
| Mouse IgG1 | NeuN | 1:500 | Millipore | MAB377 |
| Mouse IgG2a | GAD67 | 1:500 | Millipore | MAB5406 |

**Table 2B.** Secondary antibodies for interneuron analysis

| Host | Epitope | Fluorophore | Concentration | Company | Order number |
| --- | --- | --- | --- | --- | --- |
| Goat | Anti-rat | Alexa Fluor 568 | 1:500 | Life technologies | A11077 |
| Goat | Anti-rabbit | Alexa Fluor 405 | 1:500 | Life technologies | A31556 |
| Goat | Anti-mouse IgG1 | Alexa Fluor 647 | 1:500 | Life technologies | A21240 |
| Goat | Anti-mouse IgG2a | Alexa Fluor 488 | 1:500 | Life technologies | A21131 |

**Table 3A.** Primary antibodies for synapse analysis

| Host | Epitope | Concentration | Company | Order number |
| --- | --- | --- | --- | --- |
| Chicken | VGAT | 1:1000 | Synaptic systems | 131006 |
| Rabbit | PV | 1:1000 | Life technologies | PA1933 |
| Guinea pig | NeuN | 1:500 | Millipore | ABN90 |
| Mouse | CB1-R | 1:1000 | Synaptic systems | 258011 |

**Table 3B.** Secondary antibodies for synapse analysis

| Host | Epitope | Fluorophore | Concentration | Company | Order number |
| --- | --- | --- | --- | --- | --- |
| Goat | Anti-chicken | Alexa Fluor 647 | 1:500 | Life technologies | A21449 |
| Goat | Anti-rabbit | Alexa Fluor 405 | 1:500 | Life technologies | A31556 |
| Goat | Anti-guinea pig | Alexa Fluor 568 | 1:500 | Life technologies | A11075 |
| Goat | Anti-mouse | Alexa Fluor 488 | 1:500 | Life technologies | A11029 |

#### Image acquisition and analysis

Images were taken with a Zeiss Confocal microscope (type LSM700). The investigator was blinded to the groups of the sections when acquiring the images and performing the quantifications. Image analysis was performed in ImageJ (National Institutes of Health USA).

#### Cell density analysis

z-stacks were acquired at 20x of all layers of the mPFC. Tile scan z-stacks (1600 x 1280 µm^2^, 2 µm steps, total of 10 µm) were acquired of the mPFC in both hemispheres of control (n=6) and SPD (n=6) rats. Antibodies staining for NeuN, Ctip2, GAD67 and PV were used. NeuN (neuronal nuclei) is a nuclear protein specific for neurons and was used as a marker to identify neurons. Expression of Ctip2 (CtBP (C-terminal binding protein) interacting protein) is restricted to L5/6 and was used to facilitate identification of the cortical layers., The GABA synthesis enzyme GAD67 (glutamate decarboxylase) and the calcium-buffering protein PV (parvalbumin) were used to identify inhibitory cells.

#### Synapse analysis

z-stacks were acquired at 63x in layer 5 of the mPFC. For each of the rat brains (Control (n=6), SPD (n=6)) z-stacks (102 x 102 µm^2^, 0.4 µm steps, total of 12 µm) were acquired in both hemispheres. Image analysis was performed semi-automatically using custom-written ImageJ macros and MATLAB scripts. NeuN was used to determine the outline of individual L5 cell somata. VGAT (vesicular GABA transporter) is expressed in all inhibitory synapses and was used as a general inhibitory synaptic marker. For each L5 cell, a maximum intensity image was constructed from 4 z-stack slices, which was median filtered and thresholded. Only synaptic puncta larger than 0.2 µm and circularity of 0.6-1.0 that were inside of the 1.5 µm band around the NeuN outline were included. PV and CB1-R puncta were only included when co-localized with VGAT.

### Data processing and statistical analyses

Statistical analyses were performed with GraphPad Prism (Software Inc.) and RStudio 1_2_5019 (R version 3.6.1, R Foundation for Statistical Computing). Normality of the data was tested with a Shapiro-Wilk test. Differences between two groups were then tested with a nonparametric Mann-Whitney-Wilcoxon test (MW), or a parametric Welch t test (T). Differences between three groups were tested with an one-way ANOVA followed up with a Tukey’s test. Behavior in the PRL was analyzed using two-way repeated measures ANOVA (with sessions as within-subjects factor and housing condition as between-subjects factor) was used for multiple comparisons followed by T-tests (with Bonferroni correction). Detailed statistical information of the figures is listed in the Statistical Table. All graphs represent the mean ± standard error of the mean (SEM) with individual data points shown in colored circles.

## Notes

### Competing Interest Statement

The authors have declared no competing interest.

### Summary of Updates

We repeated the experiments (and replicated our earlier findings), but now added an extra group of rats that was allowed to play daily for 1 hr during the deprivation period.

